# Primitive RNA-catalysis with guanine-rich oligonucleotide sequences – the case of a (GGC)_3_ nonamer

**DOI:** 10.1101/2020.05.04.075614

**Authors:** Giovanna Costanzo, Angela Cirigliano, Samanta Pino, Alessandra Giorgi, Ondrej Šedo, Zbyněk Zdráhal, Petr Stadlbauer, Jiří Šponer, Judit E. Šponer, Ernesto Di Mauro

**Affiliations:** Institute of Molecular Biology and Pathology, CNR, Piazzale A. Moro 5, 00185, Rome, Italy; Department of Ecological and Biological Sciences, Via S. Camillo de Lellis, University of Tuscia, 01100, Viterbo, Italy; Dipartimento di Scienze Biochimiche, “Sapienza” Università di Roma, Piazzale Aldo Moro 5, 00185 Rome, Italy; Central European Institute of Technology, Masaryk University, Campus Bohunice, Kamenice 5, 62500 Brno, Czech Republic; Institute of Biophysics of the Czech Academy of Sciences, Královopolská 135, 61265 Brno, Czech Republic

**Keywords:** RNA, catalytic activity, sequence selectivity

## Abstract

A cornerstone of molecular evolution leading to the emergence of life on our planet is associated with appearance of the first catalytic RNA molecules. A question remains regarding the nature of the simplest catalytic centers that could mediate the chemistry needed for RNA-catalysis. In the current paper we provide a new example supporting our previously suggested model proposing that transiently formed open loop geometries could serve as temporary catalytic sites in the most ancient short oligonucleotides. In particular, using two independent detection techniques, PAGE and MALDI-ToF analysis, we show that prolonged thermal treatment of a 5’-phosphorylated (GGC)_3_ sequence at weakly acidic or neutral pH in the presence of tris(hydroxymethyl)aminomethane, produces a species characterized by a (GGC)_3_G stoichiometry, which is compatible with the cleavage-terminal recombination chemistry suggested in our previous studies. Our new findings are complemented by microsecond-scale molecular dynamics simulations, showing that (GGC)_3_ dimers readily sample transient potentially catalytic geometries compatible with the experimentally observed terminal recombination chemistry.

## Introduction

While modern RNA-catalysis utilizes mechanisms that are optimized in order to be less sequence specific, in the prebiotic world the situation was most likely different. Our studies on the polymerization efficiency of cyclic nucleotide precursors show that 3’,5’ cyclic GMP has a better polymerization potential than other 3’,5’ cyclic nucleotides (Costanzo et al. 2009; Costanzo et al. 2012; Šponer et al. 2015; Šponer et al. 2017). For example, drying of 3’,5’ cyclic GMP at slightly elevated temperatures commonly leads to formation of oligoG sequences with length of ca. 20 nucleotides on a timescale of several hours (Morasch et al. 2014; Šponer et al. 2015). In order to polymerize 3’,5’ cyclic AMPs higher temperatures and prolonged heat treatment are needed and, in spite of this, the maximum length of oligomers attained is ca. 4 nucleotides only (Costanzo et al. 2016). 3’,5’ cyclic CMPs dimerize and trimerize exclusively when ensuring special conditions, i.e. radical initiation (Costanzo et al. 2017) while 3’,5’ cyclic UMPs do not polymerize at all to our best knowledge (Šponer et al. 2017). All this indicates that in the prebiotic world there was an evolutionary pressure (most likely the ability of the nucleotide monomers to aggregate or crystallize, see Šponer et al. 2017) favoring G-rich sequences as the first RNA molecules, helping then in incorporation of the other nucleotides.

We hypothesized that the next level of chemical complexity might arise due to primitive forms of RNA catalysis related to WC-complementary oligoG and oligoC sequences. Our observations in Pino et al. 2013 and Stadlbauer et al. 2015 supported this proposal by suggesting that upon heat-treatment (at 60-80 °C) sufficiently (>9 nt) long oligoG and oligoC sequences are able to react with each other resulting in the transfer of the 5’-terminal nucleotide of the donor strand to the 3’-end of the acceptor. The experimentally observed reactivity was associated with minor populated states within ensembles of self-complementary oligoG and oligoC sequences in which the register of oligonucleotide strand pairing is shifted by 3-4 nucleotides enabling transient formation of catalytically active geometries. We assumed that these temporarily formed catalytic centers enabled a two-step, cleavage-terminal recombination chemistry according to a tentative mechanism explained in the left panel of Figure 1). In Stadlbauer et al. 2015, based on molecular dynamics (MD) simulations, we propose that unpaired G_4_ and C_4_ overhangs may easily fold into open tetraloops which are especially well-suited for this purpose. We note that the prerequisite for the existence of this temporarily formed geometries is that the overhangs are connected to sufficiently long WC-paired sequences (consisting of ca. six base pairs, Stadlbauer et al. 2015).

**Figure 1.**
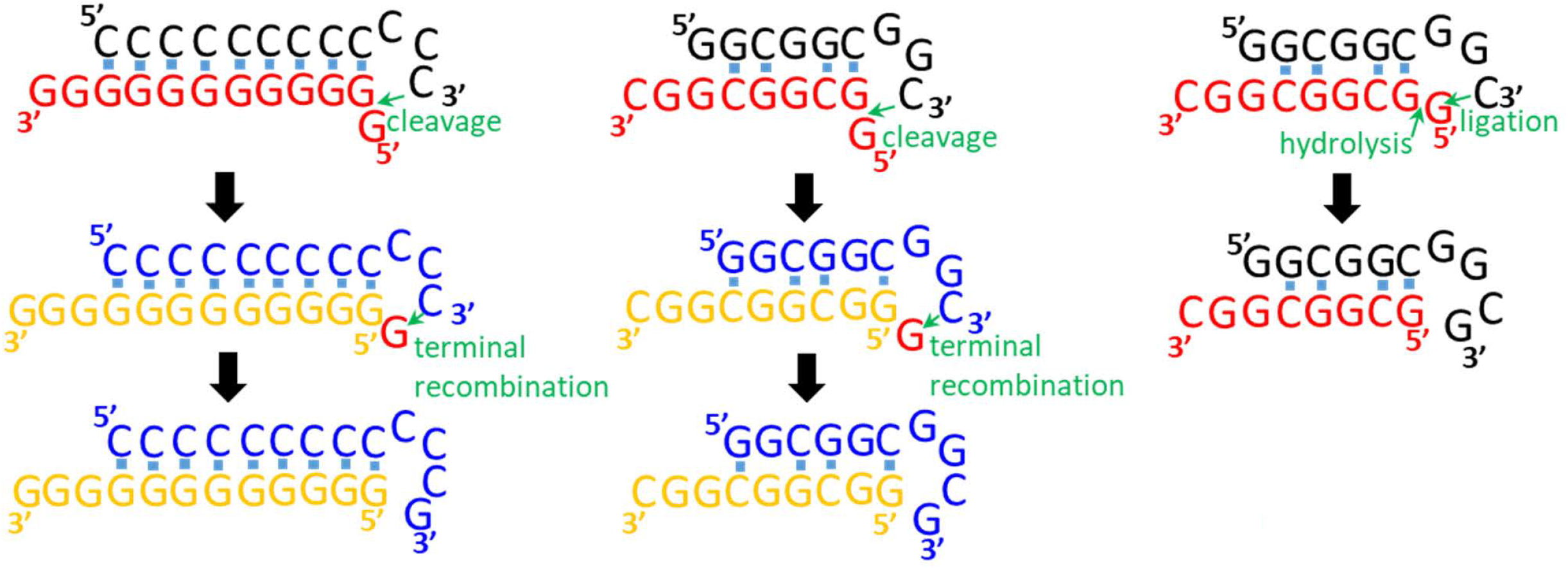
Terminal recombination chemistry proposed in our previous studies (Pino et al. 2013; Stadlbauer et al. 2015) and in the current paper. On the left: a mechanism depicting formation of a C12G oligonucleotide. The C12 acceptor strand (black) cleaves the 5’-terminal nucleotide of the donor G12 strand (red). This is followed by a terminal recombination reaction between the 3’-end of another C12 acceptor (blue) and the cleaved G nucleotide (red) produced in the preceding cleavage reaction. In the middle: the analogous chemistry with a (GGC)_3_ sequence. On the right: simplified, hydrolysis assisted mechanism. In this case the temporarily formed open tetraloop stabilizes the hydrolytically cleaved 5’-terminal nucleotide of the donor strand (red) in the vicinity of the 3’-end of the acceptor strand (black) enabling a terminal recombination reaction to take place inside the catalytic loop structure.

In the current paper we show that also mixed G- and C-containing sequences may exhibit similar catalytic properties. Our data acquired for a 5’-phosphorylated (GGC)_3_ sequence with two independent experimental techniques (PAGE and MALDI-ToF analyses) demonstrate spontaneous formation of a (GGC)_3_G decamer at 5≤pH≤7 in the presence of tris(hydroxymethyl)aminomethane (Tris). From a prebiotic chemistry perspective, this observation suggests that transient (short-lived) folded overhang structures in cooperation with amino-alcohols could play a decisive role at the advent of RNA-catalysis on our planet. Rarely populated (transient) but highly reactive conformations are suggested to play role even in modern biochemical systems (Górecka et al. 2019).

## Results and Discussion

Denaturing PAGE analysis of a 5’-phosphorylated (GGC)_3_ sequence is shown in Figure 2. We have observed weak, but well-defined new bands above that of a (GGC)_3_ sequence after prolonged (5 and 7 hours) sample treatment at 75 and 80 °C, at weakly acidic or neutral pH in the presence of Tris. As Figure 2, panel B illustrates, one can clearly exclude the possibility that the new bands are due to gel overloading.

**Figure 2.**
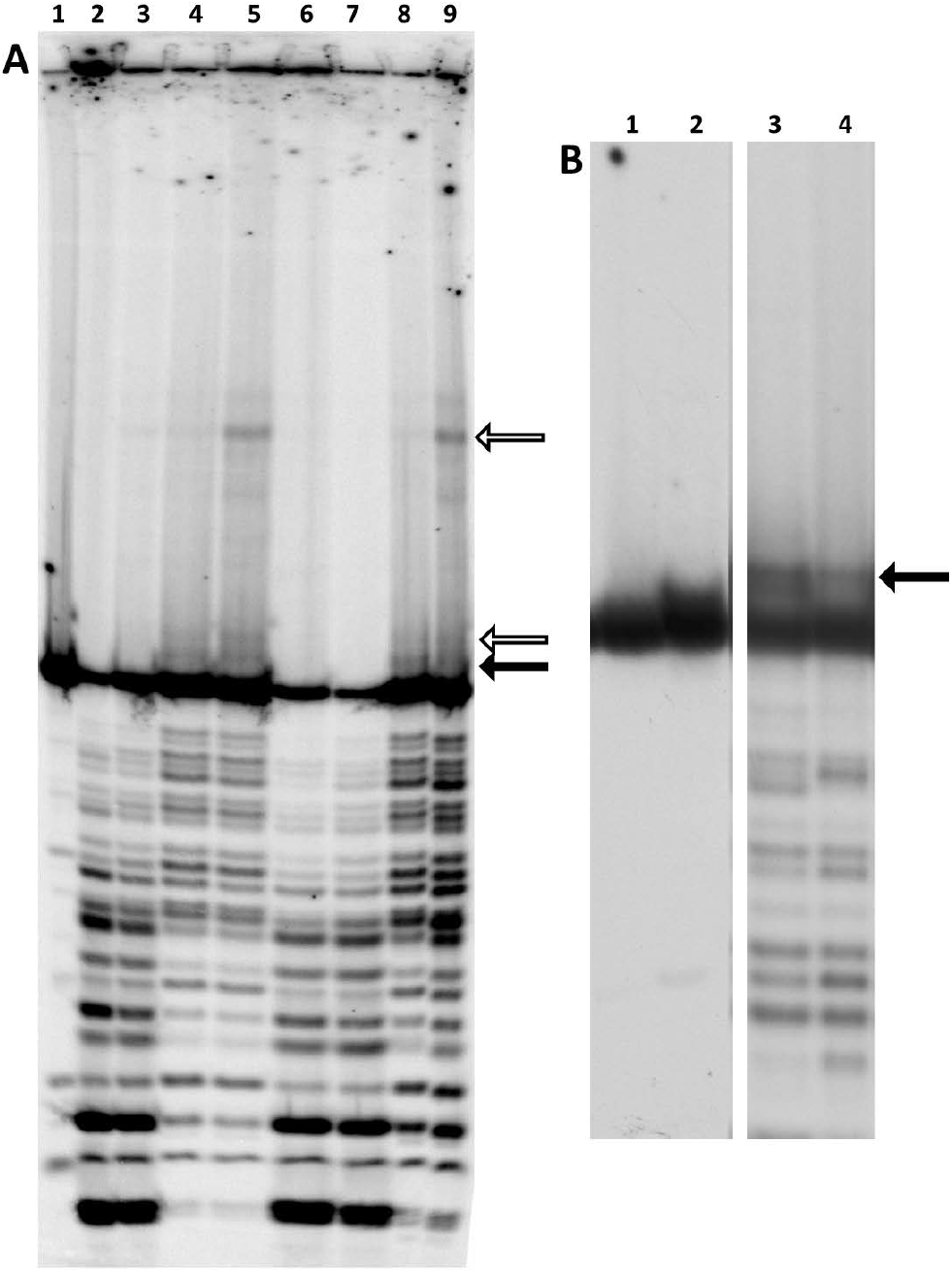
PAGE analysis of a 5’-phosphorylated (GGC)_3_ sequence treated for 5 and 7 hours in water and in Tris HCl. Panel A: Lane 1: untreated material; lanes 2-9: 5 ng of 5’-^32^P labelled (GGC)_3_ treated in water (pH=6.4) or in 10 mM Tris HCl (pH=5) under the following conditions. Lanes 2-3: 75 °C, in water, 5 or 7 hours, respectively. Lanes 4-5: 75 °C, in TrisHCl, 5 or 7 hours, respectively. Lanes 6-7: 80 °C, in water, 5 or 7 hours, respectively. Lanes 8-9: 80 °C, in TrisHCl, 5 or 7 hours, respectively. Panel B: Left: 5 (lane 1) and 15 (lane 2) ng of untreated material loaded on the gel. Right: 5ng of the treated material loaded on the gel. Sample treatment: in 10mM TrisHCl (pH=7.0) at 80 °C for 7 (lane 3) and 18 (lane 4) hours. Positions of new bands resulting from sample treatment are indicated by arrows. Black arrows: corresponding signal also found by MALDI analysis. White arrows: in lack of corresponding MALDI MS signal identification of the band is ambiguous.

We have carried out a densitometric analysis, using the positions of degradation bands of the non-treated sample for size attribution. This analysis suggested that the new bands may belong to (GGC)_3_ oligomers extended by 1 or 2 guanosine-monophophate units, i.e. (GGC)_3_G and (GGC)_3_GG. In order to unambiguously identify the new species formed we have performed a MALDI-TOF MS analysis, which has confirmed that the sample treatment leads to a new band shifted by +345 Da with respect to that of the untreated material, and might indeed belong to a (GGC)_3_G sequence (see Figure 3). Let us note, that the corresponding terminal recombination product has not been observed when the analogous (GCC)3 sequence was tested in the same conditions (data not shown, see below for explanation).

**Figure 3.**
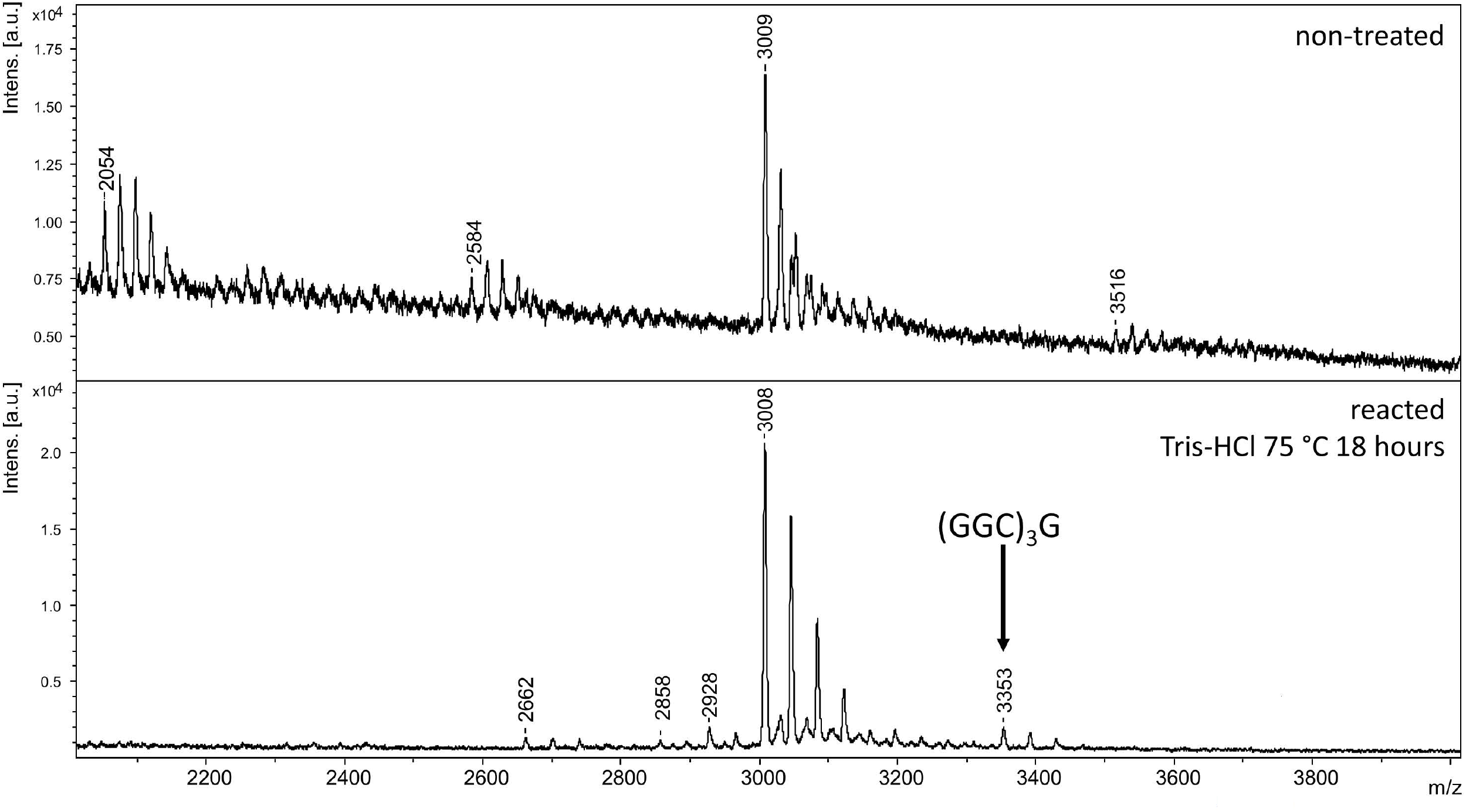
MALDI-TOF mass spectra (linear positive ion detection mode) of a (GGC)_3_ oligomer treated for 18 hours at 75 °C, pH=5.2 in the presence of 1 mM Tris HCl. After sample treatment a new signal is observed (indicated with an arrow) shifted by +345 Da with respect to the main signal at 3008, corresponding to the (GGC)_3_ oligomer. Unlabeled signals correspond to adducts with alkali metals. Equivalent signals were detected in the reflectron positive ion detection mode (not shown).

Formation of the (GGC)_3_G sequence via reaction of two (GGC)_3_ sequences can also be interpreted in terms of the previously published (Pino et al. 2013; Stadlbauer et al. 2015) two-step, cleavage - terminal recombination chemistry (cf. the left and middle panels of Figure 1). Molecular dynamics simulations provide further indication that rarely populated geometries might be responsible also for the catalytic activity observed in case of the newly-studied (GGC)_3_ sequence. We have built up a model for a (GGC)_3_-duplex (see Figure 4), assuming imperfect base-pairing with strand slippage. MD simulations reveal that within the simulation time scale the studied system readily samples geometries compatible with the cleavage reaction (see Figure 5), if the simulation is initiated from a starting geometry that is not locked by extensive intramolecular interactions (starting structures **1** and **2**, see Figure 4). Starting from conformation **3**, the reactive configuration was not reached on the simulation time scale. However, this is merely a marginal sampling issue (all three starting structures would provide identical results in the limits of infinite sampling, see Šponer et al. 2018). Note that compared to the experiments the MD time scale is very short, so there is no need to see large population of the relevant geometries. The suggested catalytic-like conformation is not a single specific loop conformation, but rather a broad ensemble of various loop shapes.

**Figure 4.**
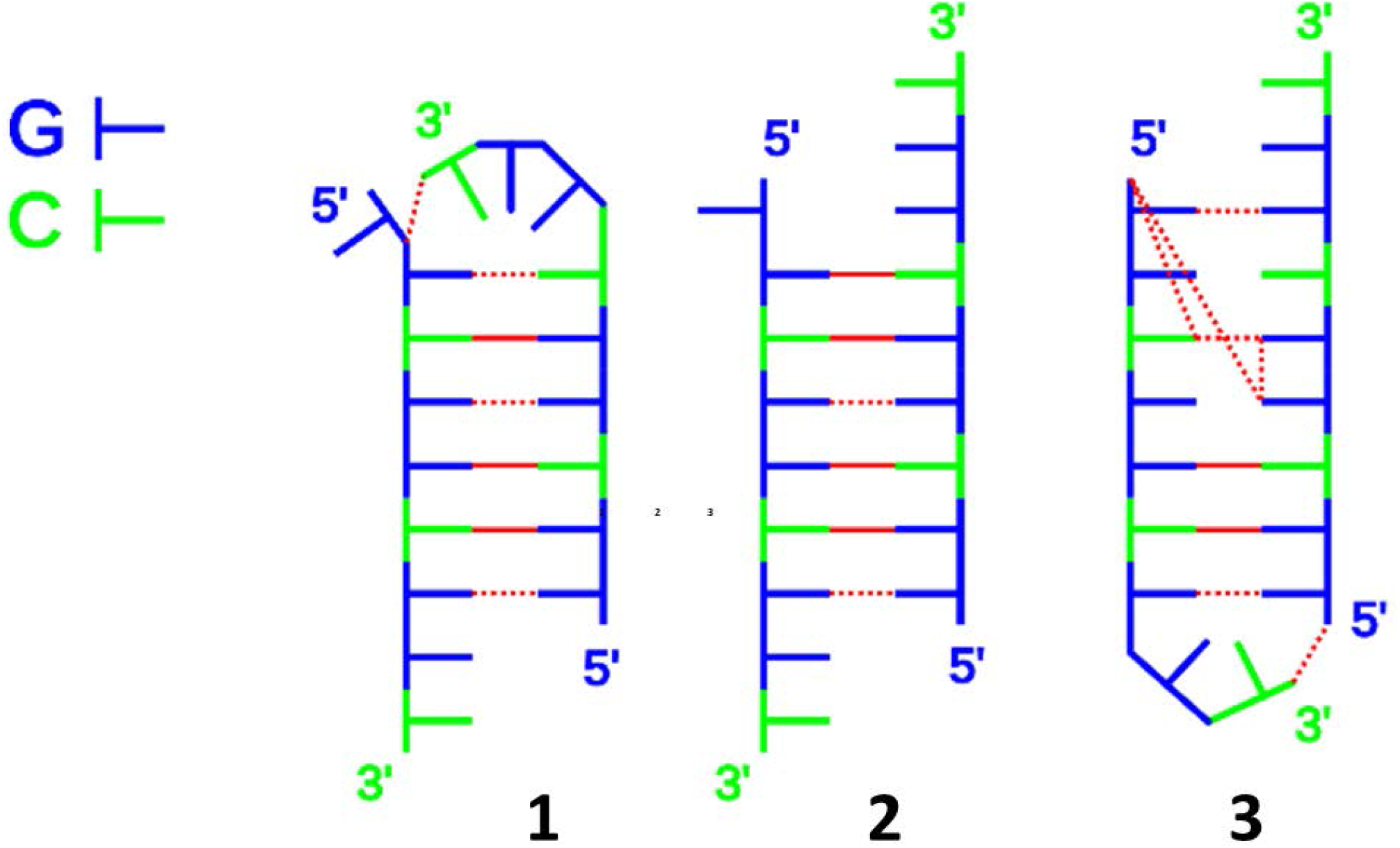
Sketch of starting geometries used for MD simulations of the model dimer formed by two slipped (GGC)_3_ strands. Solid red lines indicate canonical Watson-Crick base pairing, dotted lines stand for other hydrogen bonds.

**Figure 5.**
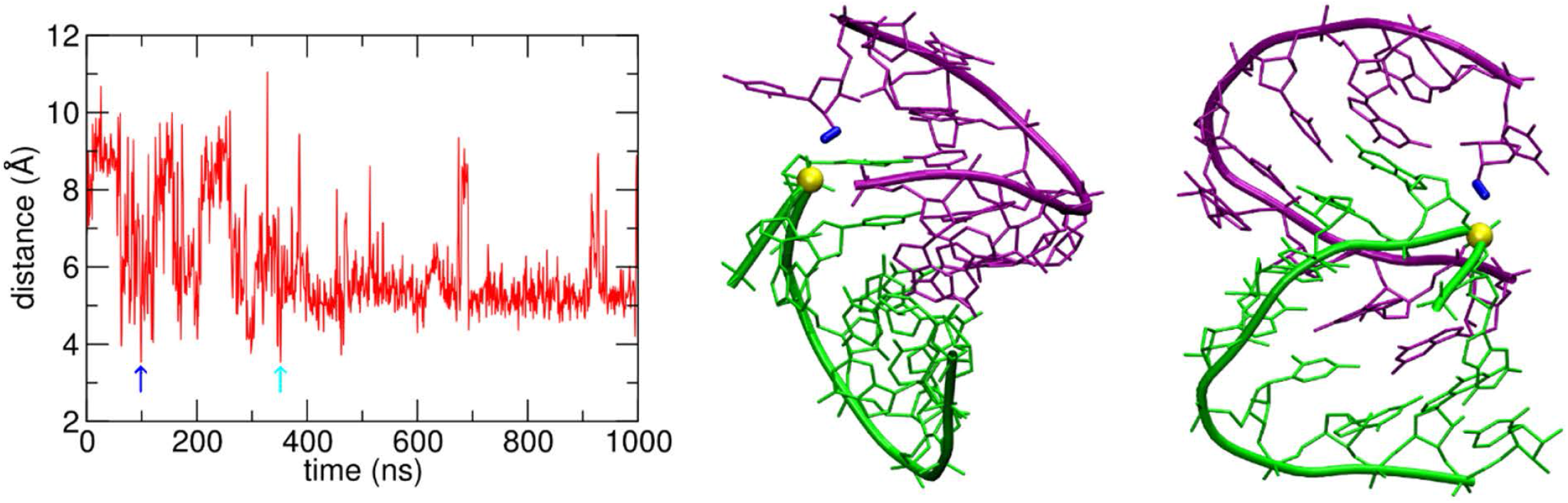
Left: Time development of the distance between the 2’-OH group of the 3’-terminal cytidine to the phosphorus atom of the subterminal phosphate group in the simulated (GGC)_3_ dimer (simulation from starting structure **2** in Figure 4 with Na^+^ counterions). The blue and cyan arrows indicate instances when the distance was shorter than 3.7 Å and accompanied with formation of H-bond to the phosphate group. Right: two side views of the potentially reactive conformation of the (GGC)_3_ dimer, at time indicated by a blue arrow in the graph on the left. Individual strands are green and purple. In this snapshot the 3’-terminal OH group (blue) of the acceptor strand contacts the 5’-subterminal phosphorus (yellow) of the donor strand. The conformation adopts a triloop structure. While MD simulations per se cannot be used to asses reaction mechanisms they can reveal intrinsic flexibility of the RNA to move towards catalytically potent geometries.

We have then considered two possible dimers formed by the (GCC)3 sequence (see Figure 6) with starting structures having equal number of H-bonds as in model **2** of Figure 4. Both duplexes have decisively melted in the course of simulations. This is in agreement with the findings by Sobczak et al. 2010 indicating that stability of GCC-based RNA duplexes is noticeably lower than those of GGC trinucleotide repeats. In other words, the double stranded core in the studied dimers of (GCC)3 sequences is too short to stabilize the slipped geometries necessary for overhang formation and for the associated ligation and terminal recombination reactions. This might explain the lack of reactivity in our experiments performed with (GCC)3 sequences. In summary, MD indicates that within the context of (GGC)_3_ strand-slipped duplex the overhang readily adopts catalytic-like conformations while analogous (GCC)3 duplexes are visibly less stable.

Our mechanistic proposal has been questioned by a recent paper (Smail et al. 2019). We would like to call attention on several key differences between our study and those reported in Smail et al. 2019. (i) In contrast to our work, the analysis performed in Smail et al. 2019 did not target the terminal recombination product directly, but rather the ligation product, which cannot be unambiguously identified on the gel shown in Figure 4 of their Supplementary information, because the denaturation is apparently not perfect and the oligomers are partially degraded. Further, the authors used entirely different sequences treated under entirely different conditions in their analysis. The gel they present is clearly contaminated by an excess of the unincorporated ^32^P label that was not removed prior to loading the analyzed material on the gel resulting in a strong background that makes the detection of the terminal recombination product impossible.

In our experiments, the terminal recombination product is observed exclusively in the presence of added Tris, which indicates that Tris cations may actively participate in the transphosphorylation chemistry. Let us note that the reaction requires heating at 75-80 °C because transphosphorylation reactions on a non-activated phosphate group are thermodynamically weakly unfavored (Staroseletz et al. 2018). With regard to kinetics, the obviously relatively high activation energy for the attack of the hydroxyl group on the phosphorus might be reduced by interaction with Tris cations. As shown in Figure 7, aminoalcohols could decrease the activation energy of the reaction by establishing more than one H-bond with the phosphate moiety that enhances the partial positive charge (and, thus, the electrophilic character) of the phosphorus atom.

The panels on the left and in the middle of Figure 1 show the originally proposed mechanism involving two acceptor oligonucleotide strands (orange and red in the Figure). On the right panel we propose a simpler, hydrolysis-assisted mechanism that could also account for the experimentally observed chemistry. It is in principle not possible to experimentally distinguish between the mechanisms involving one or two acceptor strand(s). Nevertheless, it is very unlikely that without the assistance of temporarily formed well-defined catalytic geometries, like that of tri- or tetraloops, the terminal recombination step of the hydrolysis-assisted mechanism could operate.

The remarkable difference between the behavior of the G-rich (GGC)_3_ and C-rich (GCC)3 sequences in the above described experiments suggests that at the origin of life sequence selectivity could play a decisive role. This observation is supported by earlier experimental data. For example, oligomerization of imidazole-activated guanosine-monophosphates over an oligoC template is well-known (Inoue and Orgel 1982), whereas the reverse reaction involving imidazole-activated cytidine monophosphate and an oligoG template has not been demonstrated yet (Wu and Orgel 1992). As mentioned above, cyclic nucleotide monomers remarkably differ in their propensity to oligomerize in a template-free fashion (Šponer et al. 2017). The data presented here further support the finding that guanosine-monophosphates and guanine-rich oligonucleotides, due to the distinct ability of guanine to form versatile H-bonded and stacked structures, are hot candidates to support those primitive processes that could contribute to the emergence of the catalytic function of RNA-molecules.

## Materials

2-amino-2-hydroxymethyl propane-1,3-diol (Tris BioUltra for molecular biology) was purchased from Sigma-Aldrich (cat. no. 93362). RNA oligonucleotides were purchased from Biomers.net (Ulm, Germany), and were provided phosphorylated at the 5’ extremity and in the standard dried form. Romil-SpS (Super Purity Solvent) water was used throughout.

## Methods

### Terminal Labelling and Purification of the RNA Oligonucleotides

The oligomers (300 pmol) were labelled with [γ-^32^P] ATP. Phosphorylation was carried out by adding 1 μL of T4 Polynucleotide kinase PNK (EC 2.7.1.78, 10 U/μL, New England Biolabs, Ipswich, MA, USA; # M0201L), 2 μL of 10 × PNK buffer and 1 μL of [γ-^32^P]ATP in 20μL, followed by incubation at 37 °C for 30 minutes. The oligomers were then purified on a 16% denaturing acrylamide (19:1 acrylamide/bis acrylamide, 8 M urea) gel. After elution, the RNA was precipitated by addition of 5 μl of glycogen (Thermo Scientific 20 μg/μL in water) and 3 volumes of ethanol, kept overnight at −20°C, centrifuged, washed once with a 70% ethanol/water mixture, and dehydrated (Savant SpeedVac Concentrator, 13000 rpm, 10 min, room temperature, environmental atmospheric pressure). The pellet was suspended in H_2_O, aliquoted, and conserved at −20°C.

### Reactions of the ribo-oligonucleotides and electrophoretic analyses of the reaction products

5 ng of the oligonucleotides (specific activity typically 15000 cpm/pmol) were dehydrated (Savant SpeedVac Concentrator, 13000 rpm, 10 min, room temperature, environmental atmospheric pressure) and resuspended in 15 μl of water (Romil-SpS Super Purity Solvent, pH 6.4) or in Tris HCl at 5 ≤pH≤ 7 (precise values for each particular experiment are given in the captions of Figures 2 and 3). The reaction mixture was then incubated in the temperature range 75-80 °C for 5-18 hours (precise time and temperature data are given in the captions of Figures 2 and 3 for each particular experiment). After the reaction the samples were immediately precipitated by the addition of 75 μl of Romil-SpS water, 10 μl of 3 M sodium acetate (pH 7.5), 300 μl of 96% ethanol, and 1 μl of 20 μg/μl glycogen. After precipitation the samples were suspended in 5 μl of formamide, heated for 3 min at 60°C and loaded on a 20% denaturing polyacrylamide gel (19:1 acrylamide/bis-acrylamide, 8 M urea).

### MALDI-TOF MS

The samples were mixed with the MALDI matrix (3-hydroxypicolinic acid, 75 mg/ml in 20 mg/ml dibasic ammonium citrate:acetonitrile, 1:1 v/v mixture) in 1:4 v/v ratio. After being applied to a stainless steel sample target, the samples were analyzed in linear and reflectron positive ion detection modes. The analyses were performed with an Ultraflextreme MALDI-TOF mass spectrometer (Bruker Daltonics, Bremen, Germany).

### Molecular dynamics simulations

Starting geometries for the models shown in Figures 4 and 6 were derived directly from structures previously obtained in the simulations of analogous duplexes of oligoG and oligoC sequences (Stadlbauer et al. 2015) or they were built using the Nucleic Acid Builder (Macke and Case 1997) program. Each investigated model was solvated in a truncated octahedral box of OPC water molecules (Izadi et al. 2014) with the distance of the nucleic acid to the box border being at least 10 Å. Two different neutralization schemes were applied to each system. First, net neutralization by Na^+^ cations was used; second, net neutralization by K^+^ cations followed by addition of 0.15M KCl was applied. Joung and Cheatham ion parameters for TIP4PEw were used (Joung and Cheatham 2008). This solvation scheme is common in MD simulations. Usage of H^+^ cations would seem more appropriate in the current case but it is problematic in MD simulations. Although our solvation scheme does not match the experimental conditions, our previous study on the reactivity of duplexes formed by slipped oligoG and oligoC strands has demonstrated that the occurrence of possible reactive conformations does not depend on the type of cations present in the simulation solvent (Stadlbauer et al. 2015).

The simulations were performed using the Cornell et al. force field (Cornell et al. 1995) with the parmbsc0 (Pérez et al. 2007) and parmχOL3 (Zgarbová et al. 2011) dihedral modifications, supplemented by van der Waals parameters for phosphates (Steinbrecher et al. 2012) and gHBfix potential function improving the description of RNA hydrogen bonds (Kührová et al. 2019).

**Figure 6.**
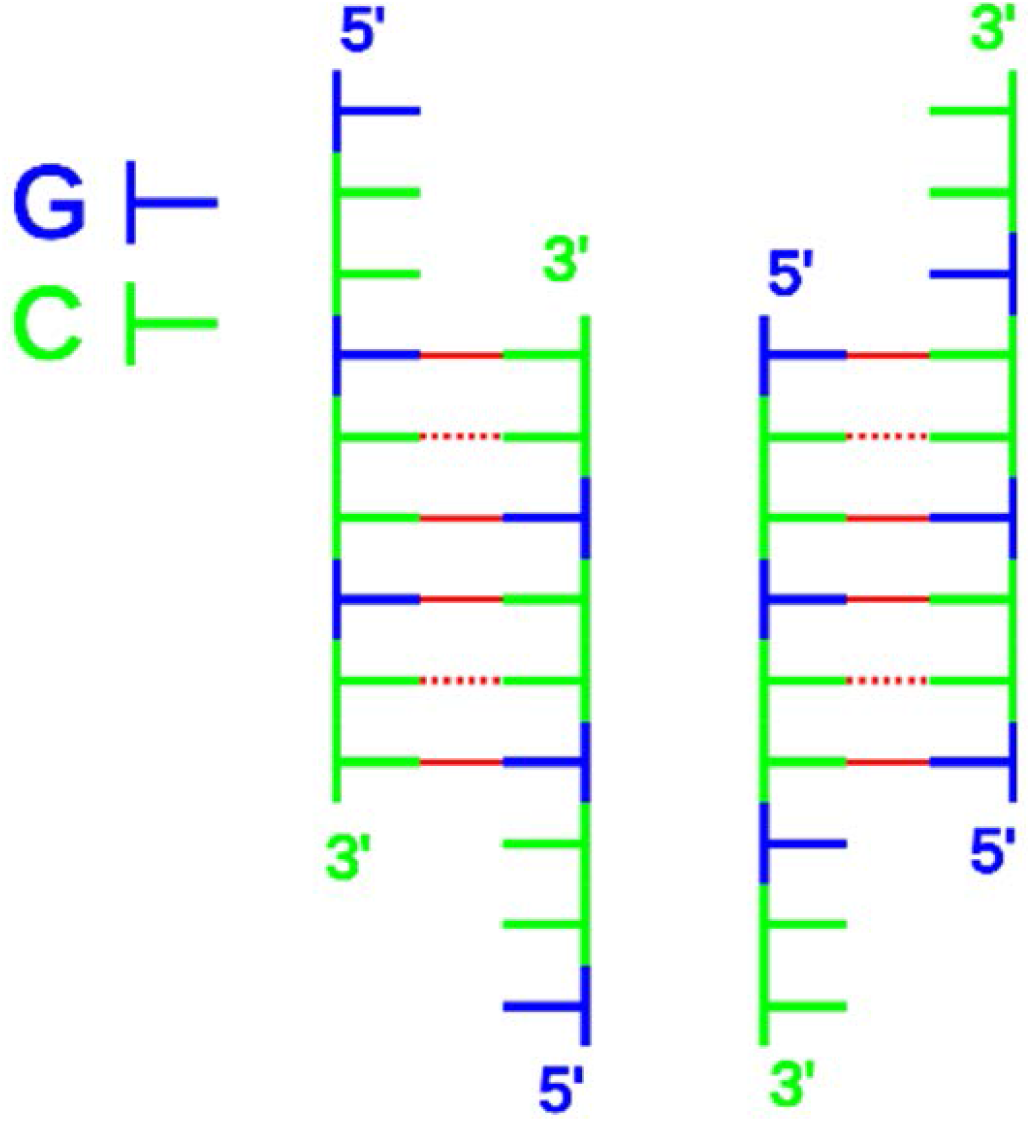
Sketch of initial geometries of the (GCC)3 dimers used for molecular dynamics simulations.

**Figure 7.**
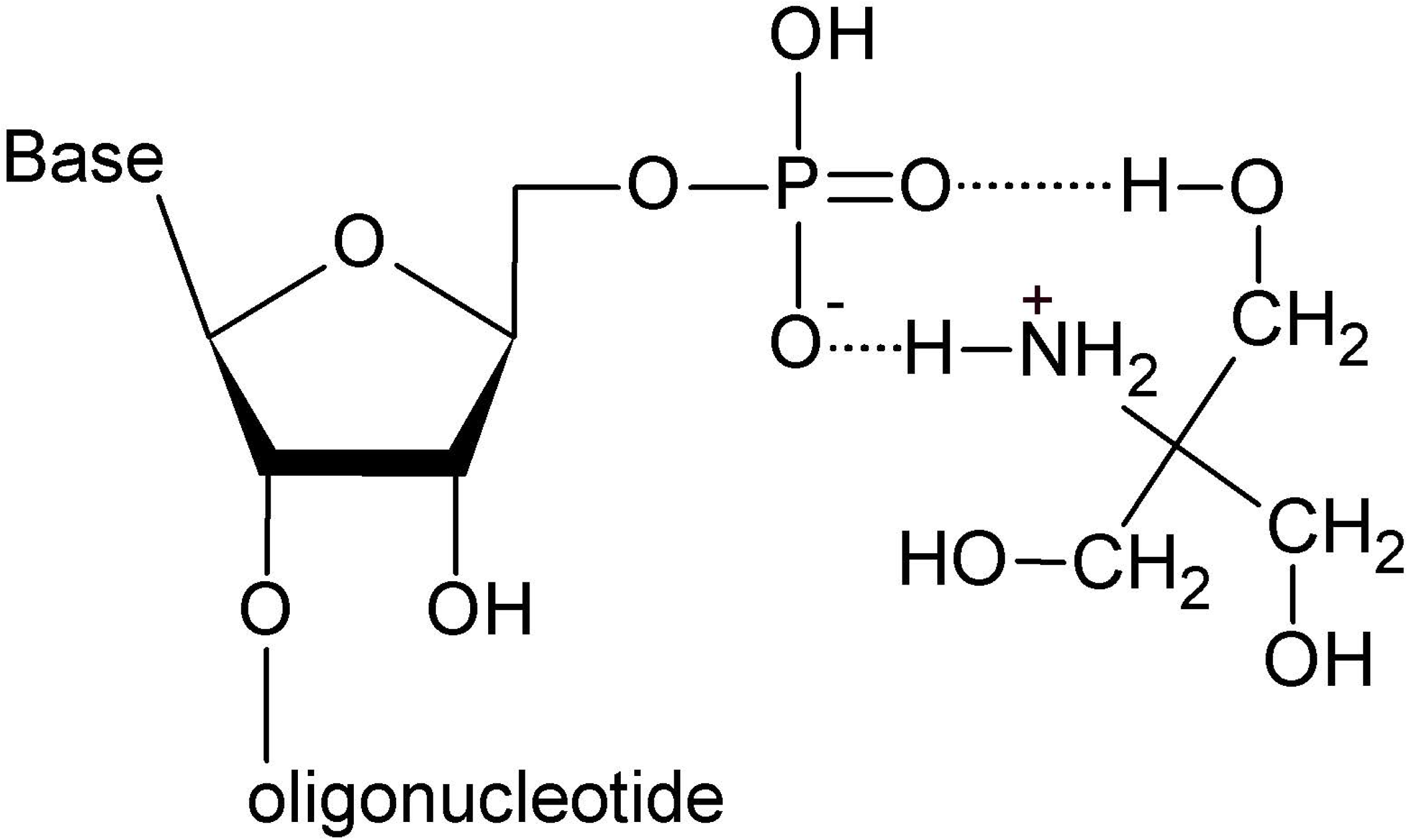
Phosphate activation by protonated amino-alcohols, shown on the example of Tris-(hydroxymethyl)aminomethane. The phosphate group may act as a multiple H-bond acceptor, while the Tris cation may serve as multiple H-bond donor. By establishing H-bonds the partial positive charge of the phosphorus increases, i.e. it becomes a better target for nucleophiles.

Starting structures were first equilibrated and then subjected to production dynamics using the standard MD protocol (Berendsen et al. 1984; Essmann et al. 1995; Ryckaert et al. 1977; Miyamoto and Kollman 1992; Hopkins et al. 2015), as described elsewhere (Islam et al. 2020; Stadlbauer et al. 2019). The calculations were done using the CUDA version of the pmemd module of AMBER 18 (Salomon-Ferrer et al. 2013; Case et al. 2018).

## Acknowledgements

This work was supported by the Italian Space Agency (ASI) project N. 2019-3-U.0, “Vita nello spazio - Origine, Presenza, Persistenza della vita nello Spazio, dalle molecole agli estremofili**”** (Space life - OPPS). We thank Dr. Aleš Kovařík for his kind help at the preparation of the samples used for MALDI MS experiments. The project CEITEC 2020 (LQ1601) and CIISB research infrastructure project LM2018127 funded by MEYS CR are gratefully acknowledged for the financial support of the MALDI MS measurements at the CEITEC Proteomics Core Facility.

## References

Berendsen HJC, Postma JPM, van Gunsteren WF, DiNola A, Haak JR. 1984. Molecular dynamics with coupling to an external bath. J Chem Phys 81: 3684–3690.

Case DA, Ben-Shalom IY, Brozell SR, Cerutti DS, Cheatham III TE, Cruzeiro VWD, Darden TA, Duke RE, Ghoreishi D, Gilson MK et al. 2018. AMBER 2018. University of California, San Francisco.

Cornell WD, Cieplak P, Bayly CI, Gould IR, Merz KM, Ferguson DM, Spellmeyer DC, Fox T, Caldwell JW, Kollman PA. 1995. A second generation force field for the simulation of proteins, nucleic acids, and organic molecules. J Am Chem Soc 117: 5179–5197.

Costanzo G, Giorgi A, Scipioni A, Timperio AM, Mancone C, Tripodi M, Kapralov M, Krasavin E, Kruse H, Sponer J et al. 2017. Nonenzymatic oligomerization of 3’,5’-cyclic CMP induced by proton and UV irradiation hints at a non-fastidious origin of RNA. ChemBioChem 18: 1535–1543.

Costanzo G, Pino S, Ciciriello F, Di Mauro E. 2009. Generation of long RNA chains in water. J Biol Chem 284: 33206–33216.

Costanzo G, Pino S, Timperio AM, Šponer JE, Šponer J, Nováková O, Šedo O, Zdráhal Z, Di Mauro E. 2016. Non-enzymatic oligomerization of 3’, 5’ cyclic AMP. PLoS One 11: e0165723.

Costanzo G, Saladino R, Botta G, Giorgi A, Scipioni A, Pino S, Di Mauro E. 2012. Generation of RNA molecules by a base-catalysed click-like reaction. ChemBioChem 13: 999–1008.

Essmann U, Perera L, Berkowitz ML, Darden T, Lee H, Pedersen LG. 1995. A smooth particle mesh Ewald method. J Chem Phys 103: 8577–8593.

Górecka KM, Krepl M, Szlachcic A, Poznański J, Šponer J, Nowotny M. 2019. RuvC uses dynamic probing of the Holliday junction to achieve sequence specificity and efficient resolution. Nat Commun 10: 4102.

Hopkins CW, Le Grand S, Walker RC, Roitberg AE. 2015. Long-time-step molecular dynamics through hydrogen mass repartitioning. J Chem Theory Comput 11: 1864–1874.

Inoue T, Orgel LE. 1982. Oligomerization of (guanosine 5’-phosphor)-2-methylimidazolide on poly(C): an RNA polymerase model. J Mol Biol 162: 201–217.

Islam B, Stadlbauer P, Vorličková M, Mergny J-L, Otyepka M, Šponer J. 2020. Stability of two-quartet G-quadruplexes and their dimers in atomistic simulations. J Chem Theory Comput, in press.

Izadi S, Anandakrishnan R, Onufriev AV. 2014. Building water models: a different approach. J Phys Chem Lett 5: 3863–3871.

Joung IS, Cheatham III TE. 2008. Determination of alkali and halide monovalent ion parameters for use in explicitly solvated biomolecular simulations. J Phys Chem B 112: 9020–9041.

Kührová P, Mlýnský V, Zgarbová M, Krepl M, Bussi G, Best RB, Otyepka M, Šponer J, Banáš P. 2019. Improving the performance of the AMBER RNA force field by tuning the hydrogen-bonding interactions. J Chem Theory Comput 15: 3288–3305.

Macke TJ, Case DA. 1997. Modeling unusual nucleic acid structures. In Molecular modeling of nucleic acids, Vol 682, pp. 379–393. American Chemical Society.

Miyamoto S, Kollman PA. 1992. Settle: An analytical version of the SHAKE and RATTLE algorithm for rigid water models. J Comput Chem 13: 952–962.

Morasch M, Mast CB, Langer JK, Schilcher P, Braun D. 2014. Dry polymerization of 3 ‘, 5 ‘cyclic GMP to long strands of RNA. ChemBioChem 15: 879–883.

Pérez A, Marchán I, Svozil D, Šponer J, Cheatham III TE, Laughton CA, Orozco M. 2007. Refinement of the AMBER force field for nucleic acids: improving the description of α/γ conformers. Biophys J 92: 3817–3829.

Pino S, Costanzo G, Giorgi A, Šponer J, Šponer JE, Di Mauro E. 2013. Ribozyme activity of RNA nonenzymatically polymerized from 3’,5’-cyclic GMP. Entropy 15: 5362–5383.

Ryckaert J-P, Ciccotti G, Berendsen HJC. 1977. Numerical integration of the Cartesian equations of motion of a system with constraints: molecular dynamics of n-alkanes. J Comput Phys 23: 327–341.

Salomon-Ferrer R, Götz AW, Poole D, Le Grand S, Walker RC. 2013. Routine microsecond molecular dynamics simulations with AMBER on GPUs. 2. Explicit solvent particle mesh Ewald. J Chem Theory Comput 9: 3878–3888.

Smail BA, Clifton BE, Mizuuchi R, Lehman N. 2019. Spontaneous advent of genetic diversity in RNA populations through multiple recombination mechanisms. RNA 25: 453–464.

Sobczak K, Michlewski G, de Mezer M, Kierzek E, Krol J, Olejniczak M, Kierzek R, Krzyzosiak WJ. 2010. Structural diversity of triplet repeat RNAs. J Biol Chem 285: 12755–12764.

Šponer J, Bussi G, Krepl M, Banáš P, Bottaro S, Cunha RA, Gil-Ley A, Pinamonti G, Poblete S, Jurečka P et al. 2018. RNA structural dynamics as captured by molecular simulations: a comprehensive overview. Chem Rev 118: 4177–4338.

Šponer JE, Šponer J, Di Mauro E. 2017. New evolutionary insights into the non-enzymatic origin of RNA oligomers. Wiley Interdiscip Rev RNA 8: e1400.

Šponer JE, Šponer J, Giorgi A, Di Mauro E, Pino S, Costanzo G. 2015. Untemplated nonenzymatic polymerization of 3 ‘,5 ‘ cGMP: a plausible route to 3 ‘,5 ‘-linked oligonucleotides in primordia. J Phys Chem B 119: 2979–2989.

Stadlbauer P, Kührová P, Vicherek L, Banáš P, Otyepka M, Trantírek L, Šponer J. 2019. Parallel G-triplexes and G-hairpins as potential transitory ensembles in the folding of parallel-stranded DNA G-Quadruplexes. Nucleic Acids Res 47: 7276–7293.

Stadlbauer P, Šponer J, Costanzo G, Di Mauro E, Pino S, Šponer JE. 2015. Tetraloop-like geometries could form the basis of the catalytic activity of the most ancient ribooligonucleotides. Chem Eur J 21: 3596–3604.

Staroseletz Y, Nechaev S, Bichenkova E, Bryce RA, Watson C, Vlassov V, Zenkova M. 2018. Non-enzymatic recombination of RNA: Ligation in loops. BBA - General Subjects 1862: 705–725.

Steinbrecher T, Latzer J, Case DA. 2012. Revised AMBER parameters for bioorganic phosphates. J Chem Theory Comput 8: 4405–4412.

Wu T, Orgel LE. 1992. Nonenzymic template-directed synthesis on oligodeoxycytidylate sequences in hairpin oligonucleotides. J Am Chem Soc 114: 317–322.

Zgarbová M, Otyepka M, Šponer J, Mládek A, Banáš P, Cheatham III TE, Jurečka P. 2011. Refinement of the Cornell et al. nucleic acids force field based on reference quantum chemical calculations of glycosidic torsion profiles. J Chem Theory Comput 7: 2886–2902.

